# A low-abundance class of Dicer-dependent siRNAs produced from a variety of features in *C. elegans*

**DOI:** 10.1101/2024.02.15.580610

**Authors:** Thiago L. Knittel, Brooke E. Montgomery, Alex J. Tate, Ennis W. Deihl, Anastasia S. Nawrocki, Frederic J. Hoerndli, Taiowa A. Montgomery

**Affiliations:** Department of Biology, Colorado State University, Fort Collins, CO 80523, USA; Cell and Molecular Biology Program, Colorado State University, Fort Collins, CO 80523, USA; Department of Biomedical Science, Colorado State University, Fort Collins, Colorado 80523, USA

**Author notes:** Equal contribution (Thiago L. Knittel, Brooke E. Montgomery).

## Abstract

Canonical small interfering RNAs (siRNAs) are processed from double-stranded RNA (dsRNA) by the endoribonuclease Dicer. siRNAs are found in plants, animals, and some fungi where they associate with Argonautes to direct RNA silencing. In *Caenorhabditis elegans*, some endogenous small RNAs, such as 22G-RNAs and 26G-RNAs, share certain attributes with canonical siRNAs but exhibit unique characteristics known only to occur in nematodes. For instance, 22G-RNAs do not originate from dsRNA and are not processed by Dicer, whereas 26G-RNAs require Dicer but lack the typical duplex intermediate with symmetrical 3’-overhangs and are produced only antisense to their mRNA templates. To identify canonical siRNAs in *C. elegans*, we first characterized the siRNAs produced from exogenous dsRNA. As predicted based on earlier studies, exogenous dsRNA is processed into ∼23-nt duplexes with 2-4-nt 3’-overhangs, ultimately yielding siRNAs devoid of 5’ G-containing sequences that bind with high affinity to the Argonaute RDE-1. Leveraging these characteristics, we searched for their endogenous counterparts and identified thousands of endogenous loci representing dozens of unique elements that give rise to mostly low to moderate levels of siRNAs, called 23H-RNAs. These loci include repetitive elements, alleged coding genes, pseudogenes, non-coding RNAs, and unannotated features, many of which adopt hairpin structures reminiscent of the hpRNA/RNA interference (RNAi) pathway in flies and mice. Our results expand the known repertoire of *C. elegans* small RNAs and demonstrate that key features of the endogenous siRNA pathway are relatively unchanged in animals.

## INTRODUCTION

siRNAs are ∼21-24-nt long non-coding RNA produced through cleavage of dsRNA by the endoribonuclease Dicer (Hamilton and Baulcombe 1999; Hammond et al. 2000; Zamore et al. 2000; Bernstein et al. 2001; Elbashir et al. 2001; Grishok et al. 2001; Ketting et al. 2001; Knight and Bass 2001). Argonaute proteins associate with siRNA duplexes, discarding one strand and retaining the other, which then acts as a guide to direct silencing of target mRNAs (Grishok et al. 2001; Hammond et al. 2001). siRNAs were discovered for their role in silencing viruses and transgenes in plants shortly after the discovery that dsRNA triggers RNA silencing in *C. elegans*, thereby revealing the specificity factor underlying the RNAi pathway (Fire et al. 1998; Hamilton and Baulcombe 1999). In addition to their roles in silencing viruses and transgenes in plants, siRNAs also target endogenous coding genes, intergenic regions, and transposons (Llave et al. 2002; Kasschau et al. 2007). Similarly, endogenous siRNAs in mice and flies derive from a variety of features, most notably transposons, pointing to a common role for siRNAs in silencing foreign elements (Chung et al. 2008; Czech et al. 2008; Ghildiyal et al. 2008; Okamura et al. 2008a; Okamura et al. 2008b; Tam et al. 2008; Watanabe et al. 2008).

*C. elegans* contains multiple distinct classes of endogenous small RNAs, but canonical siRNAs like those found in other species have not been extensively studied. What are often classified as siRNAs in *C. elegans* lack key features characteristic of siRNAs in other species (Claycomb 2014). For example, 22G-RNAs do not require Dicer and lack a dsRNA intermediate (Aoki et al. 2007; Pak and Fire 2007; Gu et al. 2009; Claycomb 2014). 26G-RNAs are similar to canonical siRNAs in their requirement for Dicer, but these small RNAs derive from an asymmetric dsRNA intermediate structure that lacks the 2-nt, 3’-overhangs characteristic of Dicer products (Ambros et al. 2003; Ruby et al. 2006; Han et al. 2009; Conine et al. 2010; Gent et al. 2010; Vasale et al. 2010; Welker et al. 2010; Fischer et al. 2011; Blumenfeld and Jose 2016). Both 22G-RNAs and 26G-RNAs are produced antisense to their mRNA templates through the activity of RNA-dependent RNA polymerases and thus are not dependent on naturally formed dsRNA, which further distinguishes them from siRNAs in mice and flies, but is a shared feature of most siRNAs in plants and fungi (Cogoni and Macino 1999; Dalmay et al. 2000; Mourrain et al. 2000; Smardon et al. 2000; Sijen et al. 2001; Volpe et al. 2002; Aoki et al. 2007; Pak and Fire 2007; Gent et al. 2009; Gu et al. 2009; Han et al. 2009; Conine et al. 2010; Gent et al. 2010; Vasale et al. 2010).

Both exogenous RNAi and viral infection trigger a silencing response involving the production of canonical exogenous siRNAs in *C. elegans* (Grishok et al. 2001; Ketting et al. 2001; Ashe et al. 2013). Furthermore, studies exploring the interplay between RNAi pathways and RNA editing by ADAR enzymes in *C. elegans* have identified numerous endogenous small RNAs with the hallmarks of canonical siRNA. For example, they appear to be processed from a dsRNA intermediate and are 22-24-nts long (Wu et al. 2011; Reich et al. 2018; Fischer and Ruvkun 2020). These siRNAs are highly upregulated in animals with loss of function in one or both ADAR genes, presumably because the base changes introduced by RNA editing disrupt the secondary structure of dsRNA, preventing its recognition or processing by Dicer (Wu et al. 2011; Reich et al. 2018; Fischer and Ruvkun 2020).

The occurrence of endogenous small RNAs resembling canonical siRNAs in ADAR mutants suggests that endogenous siRNAs may constitute a distinct class of small RNAs largely overlooked in *C. elegans*. Presumably such siRNAs are rare in wild type *C. elegans* since they have not emerged from small RNA-seq studies characterizing small RNAs. Nevertheless, even at low abundance, they could have important roles in gene regulation and thus their identification would be an important step toward a more complete understanding of the small RNA landscape of *C. elegans*.

Here we used Argonaute co-immunoprecipitation (co-IP) combined with small RNA-seq to identify and characterize >800 distinct endogenous canonical siRNAs, called 23H-RNAs. 23H-RNAs are produced from a variety of endogenous genes, including repetitive elements, pseudogenes, ncRNAs, alleged coding genes, and unannotated regions. Many of these loci adopt hairpins similar to the hpRNA pathway in flies and mice. We find that both exogenous primary siRNAs and endogenous 23H-RNAs have a high affinity for the Argonaute RDE-1. 23H-RNA loci commonly give rise to 22G-RNAs, most of which are not dependent on *rde-1*. These loci are also typically targeted by additional primary small RNAs, most notably 26G-RNAs, indicating that multiple small RNA pathways converge on loci producing dsRNA, thereby providing a robust defense against foreign elements. Our results expand the known small RNA repertoire of *C. elegans* and provide a valuable resource for exploring unique and conserved roles for small RNAs in development and genome defense.

## RESULTS

### Molecular attributes of exogenous siRNAs in *C. elegans*

In *C. elegans*, siRNAs are processed from exogenously administered dsRNA but accumulate at very low levels compared the secondary 22G-RNAs produced downstream by RNA-dependent RNA polymerases (Sijen et al. 2001; Pak and Fire 2007). Because 22G-RNAs are produced antisense to an mRNA template, high-throughput small RNA sequencing (sRNA-seq) can be used to distinguish primary siRNAs from secondary 22G-RNAs by their alignment in the sense orientation to target mRNAs (Sijen et al. 2001; Pak and Fire 2007). Although antisense primary siRNA reads are by necessity lost with this approach, we and others have used it to characterize exogenous siRNAs as predominantly 21-24 nt with a preference for a 5’ A, C, or U (Zhang et al. 2012; Svendsen et al. 2019). However, the presence of 22-nt, 5’ G-containing small RNAs aligning to the sense strand of the mRNA in these libraries points to possible contamination from 22G-RNAs. To further refine the molecular attributes of exogenous siRNAs, we did sRNA-seq on *glp-4* mutant animals following dsRNA feeding-based RNAi against the germline-expressed gene, *pos-1* (Tabara et al. 1999a). Because *glp-4* mutants lack a germline when grown at 25°C, they express very little *pos-1* mRNA and thus have minimal capacity for secondary 22G-RNA production (Beanan and Strome 1992; Svendsen et al. 2019). Therefore, *pos-1* aligning sequences will correspond almost exclusively to the primary siRNAs produced from the exogenous dsRNA (Figure 1A). Consistent with our previous findings, primary siRNAs were predominantly 22-23-nt long and depleted of 5’ G-containing sequences (Figure 1B) (Zhang et al. 2012; Svendsen et al. 2019).

**Figure 1.**
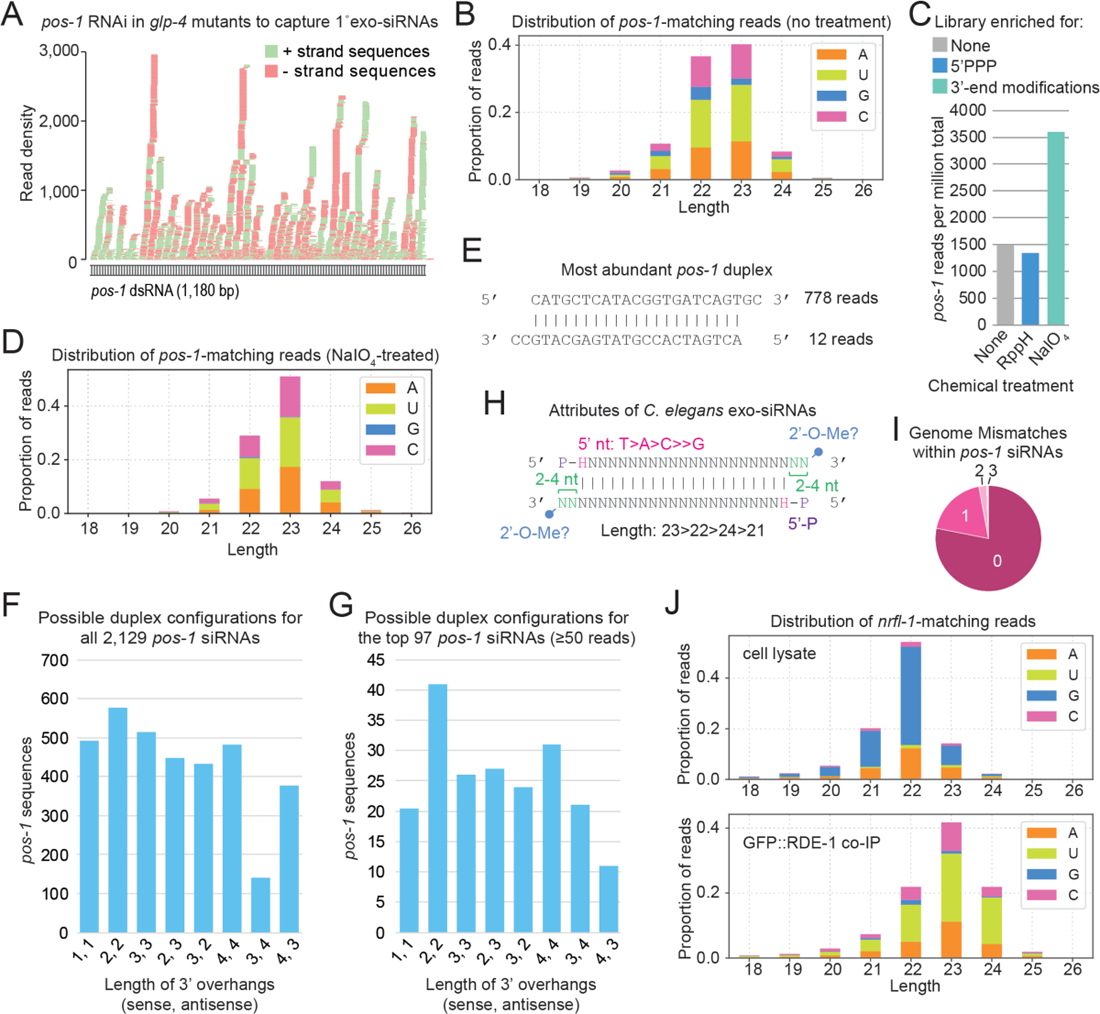
Attributes of exogenous siRNAs in *C. elegans.* (A) siRNAs produced from exogenously delivered *pos-1* dsRNA in *glp-4(bn2)* mutant worms. n=1 biological replicate. RNA from gravid adults grown at 25°C. (B) Size and 5’-nt distribution of *pos-1* siRNAs from untreated libraries. (C) *pos-1* siRNA reads from untreated libraries (none), RppH-treated libraries (reduces triphosphates), and NalO_4_-treated libraires (blocks ligation of non-3’-end-modified small RNAs). n=1 biological replicate for each treatment. RNA from gravid adults grown at 25°C. (D) Size and 5’-nt distribution of *pos-1* siRNAs from NalO_4_-treated libraries. (E) The most abundant *pos-1* siRNA duplex after *pos-1* RNAi treatment. (F) The numbers of *pos-1* siRNA sequences with a complementary strand having the indicated 3’-overhangs. Note that a single sequence could have multiple complementary strands with different 3’-overhangs. (G) As in F but limited to siRNAs with 50 or more reads. (H) Attributes of exogenous siRNAs in C. *elegans.* (I) Proportions of *pos-1* siRNA reads with 0-3 mismatches to the genome. (J) Size and 5’ nt distribution of *nrfl-1* siRNAs from cell lysates and GFP::RDE-1 co-IP libraries following *nrfl-1* RNAi. n=2 biological replicates. Data from one representative library shown. RNA from gravid adult animals.

Secondary 22G-RNAs contain a triphosphate group at their 5’ ends, whereas primary siRNAs contain a monophosphate (Pak and Fire 2007). Enzymatic treatment of small RNAs to reduce triphosphates to monophosphates prior to library construction can be used to enrich for secondary 22G-RNAs, and conversely, excluding this treatment can be used to enrich for primary siRNAs (Almeida et al. 2019).

Pretreatment of our small RNA libraries from *pos-1*-treated *glp-4*-mutants with RNA 5ʹ pyrophosphohydrolase (RppH) to reduce triphosphates to monophosphates had a negligible effect on relative *pos-1* small RNA read abundance (Figure 1C). This is consistent with our libraries having very low levels of secondary 22G-RNAs produced from *pos-1* and further confirm that primary siRNAs are monophosphorylated.

We previously showed that exogenous siRNAs are modified at their 3’ ends by the methyltransferase HENN-1 and that this modification, which is presumably 2’-O-methylation, is dependent on their association with the Argonaute RDE-1 (Svendsen et al. 2019). We observed an ∼2-fold enrichment of *pos-1*-matching reads when we pre-treated our small RNA libraries from *pos-1*-treated *glp-4*-mutants with sodium periodate, an oxidizing agent that selectively targets unmodified 3’ bases, inhibiting 3’ adapter ligation to these sequences (Figure 1C) (Seitz et al. 2008; Yu and Chen 2010; Svendsen et al. 2019). Sodium periodate treatment further enriched for 23-nt sequences and depleted 5’ G-containing sequences, likely because of loss of residual 22G-RNAs (Figure 1D).

Canonical siRNAs are processed from longer dsRNA by Dicer into ∼21-24-nt duplexes by the RNaseIII-like enzyme Dicer (Bernstein et al. 2001; Elbashir et al. 2001; Ketting et al. 2001). Like other RNaseIII cleavage products, siRNA duplexes commonly have 2-nt, 3’-overhangs (Robertson et al. 1968; Elbashir et al. 2001). Although viral siRNA duplexes commonly have 2-nt overhangs in *C. elegans,* in cell-free extracts, dsRNA is commonly processed into duplexes with 3-4-nt overhangs (Welker et al. 2011; Ashe et al. 2013).

To determine which duplex configuration is valid in exogenous RNAi, we searched our small RNA sequencing data from *pos-1* RNAi-treated animals to identify all possible duplexes. To minimize contamination of 22G-RNAs, which could confound the results, we used sRNA-seq data from libraries treated with sodium periodate, since these libraries were the most depleted of 22-nt, 5’ G-containing *pos-1* reads (Figure 1D). Even for the most abundant potential duplexes, one strand was always present at low levels, suggesting that strand stabilization is very selective (Figure 1E). Among the possible 1-4-nt 3’-end overhangs, configurations with 2-nt overhangs on either strand of the duplex were slightly more prevalent than other configurations, particularly when only including possible duplex structures in which one stand had at least 50 reads (Figures 1F-1G). Because any two complementary sequences are not necessarily from the same duplex, we cannot conclude with certainty that Dicer preferentially produces a particular 3’ overhang *in vivo* during RNAi. Furthermore, these results may be skewed by stabilization of select duplexes or single-stranded sequences downstream of dicing. Nonetheless, our data suggests that processing of dsRNA by Dicer is imprecise and yields a variety of 3’ end overhangs. In summary, we conclude that exogenous siRNAs have the following attributes: a predominant length of 23-nt, a bias against 5’ G, a 5’ monophosphate, a duplex intermediate with 2-4-nt 3’ overhangs, and a 2’-O-methyl group at their 3’ ends (Figure 1H).

### Exogenous siRNAs have very low levels of RNA editing

dsRNA is targeted by Adenosine Deaminases Acting on RNA enzymes (ADARs). ADARs function post-transcriptionally to convert A bases to inosine (Walkley and Li 2017). Inosine is read as G in high-throughput sequencing and will be reflected in the data as mismatches to the genomic sequence. Although we originally included only perfectly matching reads when analyzing our sRNA-seq data from *pos-1*-treated *glp-4*-mutants, and thus excluded reads from edited siRNAs, we repeated the analysis to include anywhere from 0-3 mismatches to the genome. Allowing for up to 3 mismatches, ∼78% of *pos-1* reads matched perfectly to the genome with 0 mismatches (Figure 1I). To determine if the mismatches present in the remaining ∼22% of aligned reads were potentially due to RNA editing, we measured the proportion of A to G mismatches (i.e. the genome nucleotide is A) for two of the most abundant siRNAs aligning to *pos-1* (Supplemental Figures S1A-S1B). Only ∼3% of siRNA reads from the most abundant siRNA analyzed had A to G mismatches (all single A to G substitution were at a single position) and none of the reads for the second most abundant siRNA had such mismatches (Supplemental Figures S1A-1B). The relatively low levels of mismatches across all reads and the very low levels of A to G mismatches in the specific siRNAs analyzed indicates that exogenous siRNAs are not typically derived from edited dsRNA. It is possible that dsRNA is less frequently edited in the cytoplasm than in the nucleus in *C. elegans* and thus exogenous dsRNA, which is unlikely to enter the nucleus, could be resistant to RNA editing (Eliad et al. 2023).

### Argonaute and Dicer underlie the molecular attributes of exogenous siRNAs

We next tested whether the 5’-nt and length profile we observed for exogenous siRNAs reflects that of the siRNAs bound to RDE-1, the primary Argonaute associated with exogenous siRNAs (Tabara et al. 1999b; Yigit et al. 2006). We co-IP’d GFP::RDE-1 from animals undergoing exogenous RNAi against both *nrfl-1* and *oma-1*, genes chosen for their mild phenotypes (Davis et al. 2022). Animals in this experiment, unlike in our *pos-1* RNAi experiments above, were able to mount a secondary 22G-RNA amplification step in response to exogenous siRNAs derived from exogenously administered *nrfl-1* and *oma-1* dsRNA. Small RNAs aligning to *nrfl-1* or *oma-1* in the cell lysate fractions were predominantly 22-nt and were strongly biased for 5’ G and were thus comprised mainly of secondary 22G-RNAs (Figure 1J; Supplemental Figure S1C). In contrast, *nrfl-1-* and *oma-1*-matching small RNA reads from the GFP::RDE-1 co-IP fractions were predominantly 23-nt and were depleted of reads containing 5’ G (Figure 1J; Supplemental Figure S1C). Thus, RDE-1-associated siRNAs mirror the cellular pool of exogenous siRNAs.

It was previously shown that *C. elegans* Dicer (DCR-1) processes dsRNA into predominantly 23-nt duplexes with variable 3’-end overhangs *in vitro*, and thus DCR-1 likely dictates the 23-nt bias and duplex structure of exogenous siRNAs (Welker et al. 2011). RDE-1, in turn, promotes 3’-end methylation, as indicated in Svendsen et al. (Svendsen et al. 2019), and may also select against 5’ G-containing small RNAs, as Argonautes are known to discriminate based on 5’-nt identity (Mi et al. 2008; Montgomery et al. 2008; Takeda et al. 2008; Frank et al. 2010; Ghildiyal et al. 2010). Thus, DCR-1 and RDE-1 are likely responsible for the molecular attributes of exogenous siRNAs described here.

Less than 1% of reads for the most abundant *nrfl-1* siRNA had A to G mismatches in either cell lysate or GFP::RDE-1 co-IP libraries (Supplemental Figure S1D). ∼2% of reads for the most abundant *oma-1* siRNA from the GFP::RDE-1 co-IP libraries had A to G mismatches, but too few reads were detected in the cell lysates for a meaningful analysis (Supplemental Figure S1E). These very low frequencies of mismatches are consistent with our earlier conclusion that exogenous siRNAs are not typically processed from edited dsRNA.

### Identification of endogenous canonical siRNAs

Using the features of exogenous siRNAs defined above, we set out to identify their endogenous counterparts. Because of the presumed rarity of endogenous siRNAs based on previous sRNA-seq experiments (Ruby et al. 2006), we sequenced small RNAs that co-IP’d with GFP::RDE-1 since RDE-1 associates with exogenous siRNAs and therefore is predicted to also associate with endogenous siRNAs (Yigit et al. 2006). We then filtered for reads 21-24-nt long, a size range that would capture most exogenous siRNAs and thus also their endogenous counterparts. We developed a computer algorithm to search these sequences for pairs that are oriented antisense to one another with 2-nt 3’ overhangs, which based on our analysis above would most effectively capture canonical siRNAs, although siRNAs with other 3’-overhang configurations would be lost. We then filtered for sequences that were represented by at least 10 reads and were enriched >2 fold in our GFP::RDE-1 co-IPs described above, relative to cell lysate input fractions (Figure 2A). We expanded our candidates to include additional RDE-1-associated sequences that aligned to loci producing small RNAs that met our selection criteria outlined in Figure 2A, but which lacked evidence for a duplex intermediate with 2-nt 3’-overhangs, since our analysis of exogenous siRNA duplexes identified other configurations as well. Using this approach, we identified 1196 candidate sequences matching perfectly to 25,701 genomic loci (reflecting the repetitive nature of many of them as described below), which like exogenous siRNAs, were depleted of 5’ G containing species (Figure 2B; Supplemental Table S1). There was a strong bias for 23-nt species, even after excluding sequences from the three most abundant siRNA-yielding loci, which made up most reads (Figures 2B-2C). We named these endogenous siRNAs 23H-RNAs because of their length and bias against 5’ G containing sequences (U, A, C = H) and to be consistent with naming of other classes of small RNAs in *C. elegans* (Ruby et al. 2006; Gu et al. 2009; Han et al. 2009; Reich et al. 2018).

**Figure 2.**
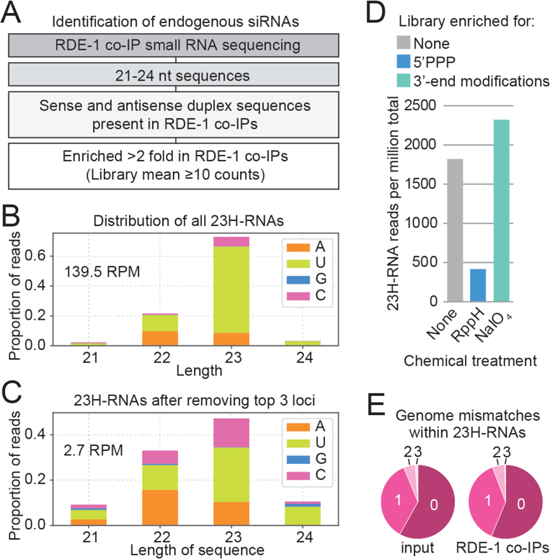
Identification of endogenous canonical siRNAs. (A) Canonical siRNA discovery flowchart. (B) Size and 5’-nt distribution of all 23H-RNAs. n=2 biological replicates. Data from one representative library shown. RNA from gravid adult animals. (C) Size and 5’-nt distribution of 23H-RNAs, as in B, after excluding the three most abundant loci. RPM, reads per million total mapped reads. (D) 23H-RNA reads from untreated libraries (none), RppH-treated libraries (reduces triphosphates), and NalO_4_-treated libraries (blocks ligation of non-3’-modified small RNAs). n=1 biological replicate. RNA from gravid adult animals. (E) Proportions of 23H-RNA reads with 0-3 mismatches to the genome. n=2 biological replicates. Data from one representative library shown. RNA from gravid adult animals.

Endogenous 23H-RNAs were depleted in wild type small RNA libraries treated with RppH due to more efficient capture of the abundant triphosphorylated 22G-RNAs, indicating that they are monophosphorylated (Figure 2D). Unlike our analysis of exogenous siRNAs, these libraries were produced from animals containing fully developed germlines which have high levels of 22G-RNAs. Hence, RppH treatment led to stronger depletion of 23H-RNAs than what we observed for exogenous siRNAs. 23H-RNAs were slightly elevated in libraries treated with sodium periodate, suggesting that they are methylated (Figure 2D) (Svendsen et al. 2019). These results are consistent with the attributes we observed for exogenous siRNAs above.

To determine if the 23H-RNAs we identified are processed from dsRNA that had been edited by ADARs, we aligned the small RNA reads from our GFP::RDE-1 co-IP and cell lysate libraries to the genome allowing for up to 3 mismatches. ∼40% of 23H-RNA reads contained 1-3 mismatches in both the cell lysate and GFP::RDE-1 co-IP libraries, which corresponds to an ∼2-fold increase relative to what we observed with exogenous siRNAs (Figures 1I and 2E). We then calculated the ratio of reads containing 1-3 genomic mismatches to total reads for each 23H-RNA, as well as for individual miRNAs for which editing is thought to be extremely rare, focusing on sequences represented by >100 total reads (Warf et al. 2012). Although the median proportion of mismatched reads to total reads was higher for 23H-RNAs than for miRNAs, there was considerable overlap in the distributions, with many 23H-RNAs having a low proportion of reads containing genomic mismatches, similar to most miRNAs (Supplemental Figure S2A; Supplemental Table S2).

To assess genomic mismatches that are more likely to be directly caused by RNA editing, we quantified the proportions of A to G mismatches for the most abundant 23H-RNA duplex, which is produced from F43E2.6. The siRNA produced from the antisense strand of the duplex, which contains 9 As in a variety of different sequence contexts, had a much higher proportion of A to G mismatches than the sense strand, which has only 3 As (Supplemental Figures S2B-S2C). ∼39% of F43E2.6 antisense siRNA reads from the cell lysates and ∼33% from GFP::RDE-1 co-IPs contained 1-3 A to G mismatches indicative of editing (Supplemental Figure S2B). Only ∼8-9% of reads corresponding to the F43E2.6 sense siRNA had A to G mismatches in either the cell lysates or GFP::RDE-1 co-IPs, suggesting that this strand with its low number of As is more resistant to editing (Supplemental Figure S2C). For each sequence, the proportion of reads containing non-A to G mismatches was similar (5-7%) and likely relates to 3’-end tailing and mutations introduced during small RNA library preparation and sequencing. Together, our suggests that there is considerable variation in the extent of editing of individual 23H-RNAs with some having very high levels of editing and others having much lower or no editing.

We next compared the abundance of 23H-RNAs in gravid adults, the stage used for their identification, with L4 stage larvae, dissected distal gonads, and embryos. Among these stages, embryos had substantially higher levels of most 23H-RNAs, although many 23H-RNAs also had elevated levels in the L4 stage relative to adults, and a small subset were elevated in dissected gonads compared to whole animals (Supplemental Figures S2D-S2F; Supplemental Table S3). Interestingly, RNA editing-enriched dsRNA regions also tend to be more highly expressed in embryos, which may reflect greater expression of transcripts forming dsRNA in embryos than in other developmental stages (Reich et al. 2018).

### 23H-RNAs are produced from a variety of features, including hpRNAs and repetitive elements

Just 3 genomic loci – F43E2.6, Y57G11C.1145, and *mir-5592* – accounted for ∼98% of 23H-RNA reads in our GFP::RDE-1 co-IP libraires (Figures 2B-2C and 3A; Supplemental Table S4). F43E2.6 is annotated as a coding gene but lacks any known function. siRNAs are processed from both strands of F43E2.6, indicating that there is overlapping transcription at the locus, although we were unable to capture sufficient RNA-seq reads to explore the extent of this overlap (Figure 3B) (Reed et al. 2020). Y57G11C.1145 is a long hairpin RNA, reminiscent of hpRNAs found in flies and mice (Figures 3A and 3C) (Czech et al. 2008; Kawamura et al. 2008; Okamura et al. 2008b; Tam et al. 2008; Watanabe et al. 2008). Indeed, many of the 23H-RNA loci adopt hairpins ranging in length from tens to hundreds of base-pairs (Figure 3C). *mir-5592* also forms a hairpin and is annotated as a miRNA but is atypical of most miRNAs in having a perfectly base-paired stem (Figure 3C). Furthermore, sequences derived from the hairpin are not enriched in co-IPs from the other miRNA-associated Argonautes – ALG-1, ALG-2, and ALG-5 (Supplemental Table S5) (Seroussi et al. 2023). We identified reads from perfectly base-paired duplexes aligning to several other miRNA loci but largely excluded them because of uncertainty in their classification. The *mir-5592* locus was retained because of the specificity of small RNAs produced from the locus for RDE-1 and its extensive base-pairing. *mir-5550*-derived small RNAs were also retained because they derive from a repetitive element which is atypical of miRNAs but common among 23H-RNAs, as described below (Supplemental Table S1). 23H-RNAs were often produced from among clusters of other smalls RNAs, with only the 23H-RNA species enriched in GFP::RDE-1 co-IPs (Figures 3B and 3D; Supplemental Figure S3A). Genes annotated as protein coding, such as F43E2.6 described above, represented ∼33% of 23H-RNA loci yielding >10 reads per million total reads (rpm) in GFP::RDE-1 co-IPs, however, it is unclear if these genes truly encode functional proteins (Figure 3E). Another ∼19% of these relatively high 23H-RNA-yielding loci are annotated as pseudogenes, a designation that may extend to many of the 23H-RNA loci annotated as coding genes (Figure 3E). However, the second largest source of 23H-RNAs was repetitive elements (Figure 3E). ∼28% of 23H-RNA sequences were derived from loci with >2 genomic copies, with ∼4% from loci with >100 copies (Figures 3F-3G). These loci were largely related to DNA transposons, such as CELE14B, DNA-8-1, PAL8A, PALTA3, PALTA4, PAL5A, and TC5A, pointing to a possible role for 23H-RNAs in transposon silencing (Figures 3A and 3G).

**Figure 3.**
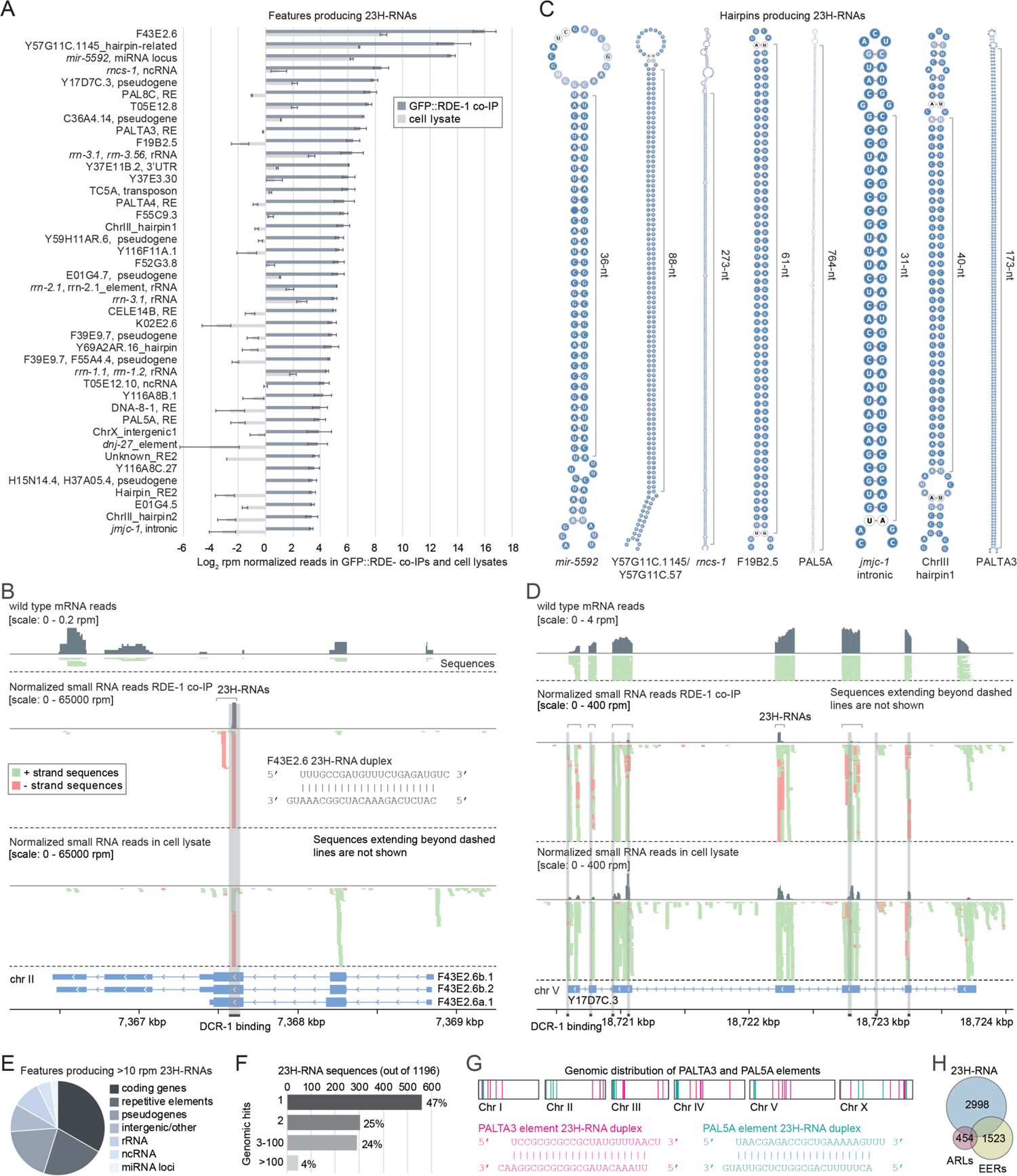
Genic and intergenic origins of 23H-RNAs. (A) Mean log_2_ rpm-normalized counts from cell lysates and GFP::RDE-1 co-IPs for features with >10 rpm in GFP::RDE-1 co-IPs. Error bars are standard deviation from the mean for 2 biological replicates. RNA from gravid adult animals. (B) mRNA reads from wild type animals and small RNA-seq reads from GFP::RDE-1 co-IPs and cell lysates plotted along the F43E2.6 23H-RNA locus. Brackets indicate 23H-RNA clusters. The sequence of the most abundant duplex is shown. The highlighted regions indicate DCR-1-binding sites based on DCR-1 PAR-CLIP data. (C) Secondary structures for several 23H-RNA loci that form hairpins. (D) As in B but for the gene Y17D7C.3. (E) The proportion of 23H-RNA loci with >10 rpm in GFP::RDE-1 co-IPs overlapping the indicated features. (F) 23H-RNA sequences categorized by number of perfectly matching genomic loci. (G) PALTA3 and PAL5A genomic distribution and the most abundant duplex from each repetitive element. (H) Overlap between 23H-RNA loci and the editing-enriched regions (EERs) identified in Reich et al. (Reich et al. 2018) and the ADAR-modulated RNA loci (ARLs) identified in Wu et al. (Wu et al. 2011). Numbers shown are for total features. 449 23H-RNAs and EERs overlap; 129 23H-RNAs and ARLs overlap; 93 EERs and ARLs overlap; 24 in common to all 3 datasets.

### Dicer’s role in 23H-RNA biogenesis

The majority of relatively high-yielding 23H-RNA loci were identified as Dicer substrates in PAR-CLIP experiments, suggesting that they are processed through Dicer cleavage as we predicted based on their duplex structure (Figures 3B and 3D; Supplemental Table S6) (Rybak-Wolf et al. 2014). We confirmed that *dcr-1* knockdown by RNAi reduced the levels of the most abundant F43E2.6 siRNA using qRT-PCR, further supporting a role for Dicer in 23H-RNA biogenesis (Supplemental Figure S3B).

Dicer’s helicase domain is required to produce a subset of endogenous small RNAs, most notably 26G-RNAs, some of which are also elevated in ADAR mutants (Welker et al. 2010; Warf et al. 2012). The helicase domain is important for processing of dsRNA containing blunt or 5’-end single-stranded overhangs but not for dsRNAs with 3’-end single stranded overhangs (Welker et al. 2011). To determine if 23H-RNAs are dependent on Dicer’s helicase domain, we reanalyzed sRNA-seq data from *dcr-1* mutants rescued with either wild type *dcr-1* or a helicase mutant form of *dcr-1* (Welker et al. 2010; Welker et al. 2011). 23H-RNAs were only modestly affected in animals containing the *dcr-1* helicase mutant, similar to some non-*dcr-1-*dependent small RNAs and in contrast to ERGO-1 class 26G-RNAs which were nearly completely lost (Supplemental Figure S3C). Interestingly, most 23H-RNA loci, including repetitive elements and even long hairpins, gave rise to a small number, often only 1 or 2, dominant duplexes (Figures 3B, 3D, and 3G; Supplemental Figure 3A). The helicase domain is thought to promote Dicer processivity and given that 23H-RNA are not dependent on this domain, it stands to reason that their biogenesis would not be processive, which may explain why so few distinct 23H-RNAs are produced from each locus (Welker et al. 2011).

### Overlap between 23H-RNAs and RNA-edited loci

We predicted that 23H-RNAs would largely overlap with other endogenous canonical siRNAs identified in studies exploring RNA editing by the ADAR enzymes since they have similar molecular attributes (Wu et al. 2011; Reich et al. 2018). To explore this, we first clustered 23H-RNA loci that were within 1,000 bp of each other, which yielded 2,998 distinct regions, which we then compared to ADAR edited regions identified previously. Of the 2,998 23H-RNA loci identified here, 454 overlapped with the 1,523 editing-enriched regions (EERs) identified in Reich et al. (Reich et al. 2018) and 129 overlapped with the ADAR-modulated RNA loci (ARLs) identified in Wu et al. (Wu et al. 2011) (Figure 3H; Supplemental Table S7). In total, we identified 2,444 23H-RNA loci that did not overlap with siRNA clusters identified in previous studies. We likely failed to identify many of the ARLs and EERs because our analysis included only animals proficient in RNA editing, and many of these loci do not produce readily detectable levels of siRNAs in wild type animals (Wu et al. 2011; Reich et al. 2018). Conversely, previous studies with a focus on RNA editing excluded siRNA clusters from sites that are not heavily edited, which presumably constitutes many of those identified here, as described above (Supplemental Figure S2A).

### RDE-1 has a high affinity for exogenous and endogenous siRNAs

Although RDE-1 associates predominantly with miRNAs, ∼14% of the total sRNA-seq reads from GFP::RDE-1 co-IPs corresponded to the 23H-RNA sequences annotated here (Figure 4A; Supplemental Table S4) (Correa et al. 2010; Svendsen et al. 2019; Seroussi et al. 2023). Despite being a minority fraction of RDE-1 associated small RNAs, there was tremendous enrichment for individual 23H-RNAs in GFP::RDE-1 co-IPs compared to most miRNAs (Figure 4B; Supplemental Table S4). Endogenous 23H-RNAs and exogenous 1° siRNAs produced from *nrfl-1* and *oma-1* were all similarly enriched and tended to be among the highest abundance small RNAs in GFP::RDE-1 co-IPs, indicating that they have a higher affinity for RDE-1 than do most miRNAs and other small RNAs (Figure 4B; Supplemental Table S4).

**Figure 4.**
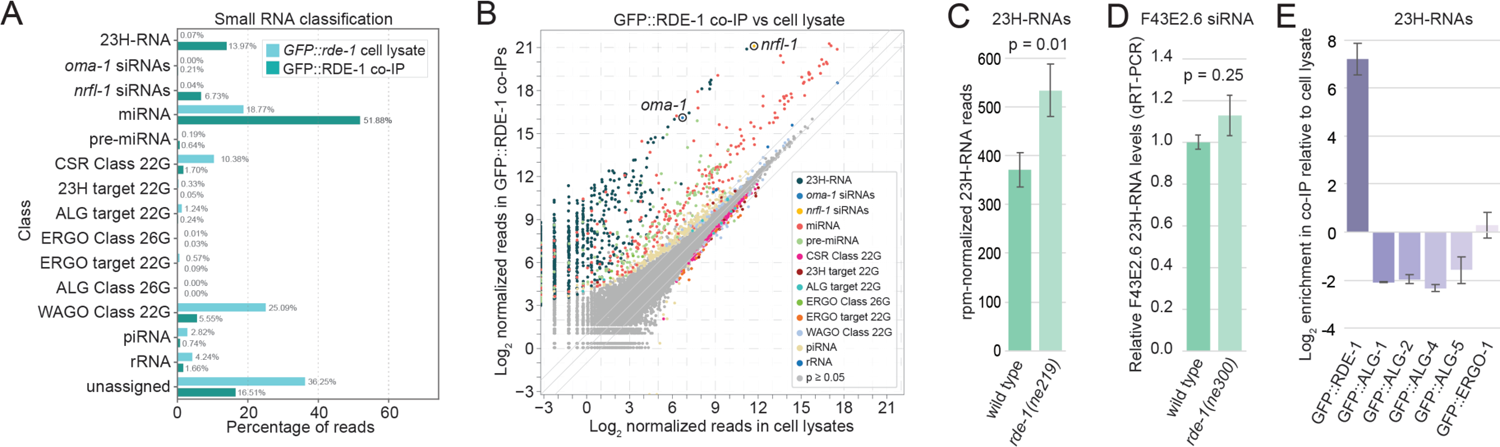
23H-RNAs have high affinity for RDE-1. (A) Percentage of reads in cell lysates and GFP::RDE-1 co-IPs corresponding to each class of small RNAs. n=2 biological replicates. RNA from gravid adult animals. (B) Each point in the scatter plot represents the average log_2_ geometric mean-normalized reads in cell lysates (x-axis) and GFP::RDE-1 co-IPs (y-axis). Exogenous siRNAs produced from *nrfl-1* and *oma-1* are circled. (C) Reads per million total mapped reads corresponding to 23H-RNAs in wild type and *rde-1(ne219)* mutants based on small RNA-seq experiments. RNA from gravid adult animals. Error bars are SD between 3 biological replicates. The p value was calculated using a two-sample t-test. (D) Relative levels of the most abundant F43E2.6 siRNA in wild type and *rde-1 (ne300)* mutants based on qRT-PCR and normalized to miR-1. RNA from gravid adult animals. Error bars are SD between 3 biological replicates. The p value was calculated using a two-sample t-test. (E) Average log_2_ enrichment of total 23H-RNA reads in the indicated Argonaute co-IPs relative to the cell lysates used in these co-IPs. Error bars are SD between 2 biological replicates. RNA from L4 stage larvae (ALG-4) or adult animals (all others).

Despite their association with RDE-1, endogenous 23H-RNAs were not depleted in *rde-1* mutants but instead appeared somewhat elevated (Figure 4C). However, RDE-1 promotes small RNA 3’ end modification, presumably 3’-2-O-methylation, which inhibits 3’-adapter ligation and can therefore confound sRNA-seq-based quantification (Svendsen et al. 2019). Using qRT-PCR we did not see significant upregulation of F43E2.6 23H-RNA levels in *rde-1* mutants, suggesting that the apparent upregulation in our sRNA-seq data may be an artifact of library preparation (Figure 4D).

Because loss of *rde-1* did not lead to a reduction in 23H-RNA levels, we tested whether these small RNAs bind to other Argonautes that might stabilize them in the absence of RDE-1. Collectively, 23H-RNAs were depleted or neutral in co-IPs of each of the other Argonautes implicated in Dicer-dependent small RNA pathways (ALG-1, ALG-2, ALG-4, ALG-5, and ERGO-1), suggesting that as a class they are preferentially bound by RDE-1 (Figure 4E) (Seroussi et al. 2023).

Nonetheless, some 23H-RNAs were enriched in one or more of the other Argonautes, indicating that they are in some instances bound by multiple Argonautes (Supplemental Figure S4; Supplemental Table S5). It is thus possible that 23H-RNAs are stabilized by these or other Argonautes in the absence of RDE-1.

### Multiple small RNA pathways converge on 23H-RNA loci

During exogenous RNAi, RDE-1 directs target mRNAs into a 2° small RNA amplification pathway to produce 22G-RNAs (Sijen et al. 2001; Pak and Fire 2007). Furthermore, RDE-1, in association with miR-243, directs a single endogenous target into the 22G-RNA pathway, leading to its silencing (Gu et al. 2009; Correa et al. 2010). To determine if endogenous 23H-RNAs also trigger 22G-RNA production, given their association with RDE-1, we searched sRNA-seq datasets from wild type animals for 22-nt, 5’ G-containing sequences, which are therefore likely to be 22G-RNAs, produced from 23H-RNA loci (Phillips et al. 2014). We detected such small RNAs at each of the 23H-RNA loci yielding >10 rpm in GFP::RDE-1 co-IPs, and consistent with their designation as 22G-RNAs, they were nearly all depleted in *mut-14 smut-1* mutants, which are defective in 22G-RNA production (Figure 5A; Supplemental Table S3) (Phillips et al. 2014). A subset of 23H-RNA loci was weakly to modestly depleted of 22G-RNAs in *rde-1* mutants, indicating that RDE-1 has a role, albeit seemingly minor, in directing 22G-RNA formation from at least some 23H-RNA targets (Figure 5B; Supplemental Table S3).

**Figure 5.**
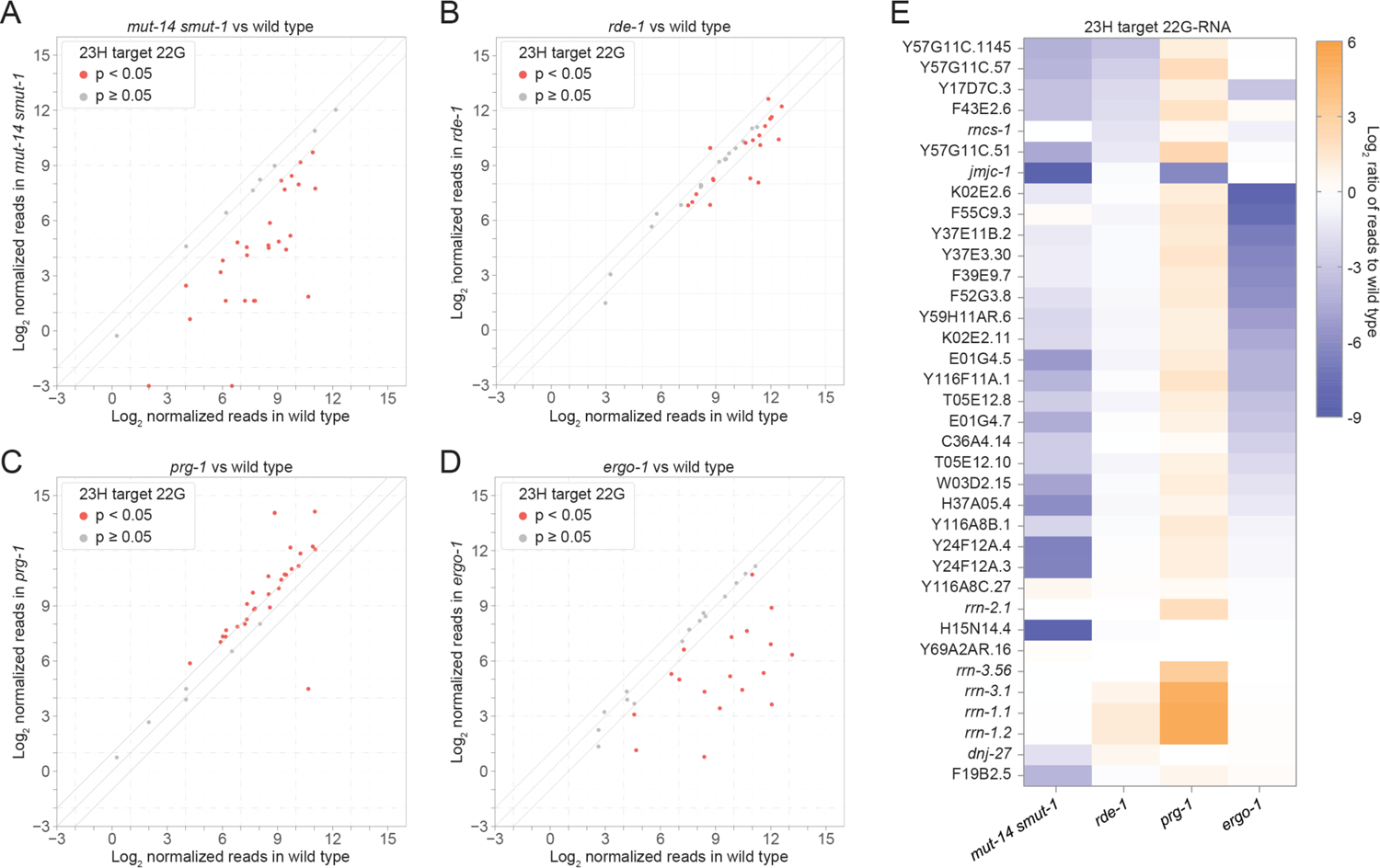
Genetic requirements for 22G-RNA formation from 23H-RNA loci. (A-D) Each point represents the average log_2_ geometric mean-normalized 22G-RNA reads for each of the top 23H-RNA loci (>10 rpm) in wild type (x-axis) and mutant (y-axis). (A) *mut-14(mg464) smut-1 (tm1301).* n=2 (wild type) or 3 *(mut-14 smut-1)* biological replicates. (B) *rde-1(ne219).* n=3 biological replicates. (C) *prg-1(n4357).* n=3 biological replicates. (D) *ergo-1(tm1860).* n=3 biological replicates. All libraries from gravid adult animals. (E) Average log_2_ ratios of 22G-RNA reads for each of the top 23H-RNA loci (>10 rpm) in the indicated mutants to wild type. Same data as in A-D.

Because many of the 23H-RNA loci were not substantially depleted of 22G-RNAs in *rde-1* mutants, we hypothesized that these loci are targeted by additional small RNA pathways. In *C. elegans*, piRNAs also direct their mRNA targets into the 22G-RNA pathway (Bagijn et al. 2012; Lee et al. 2012). In *prg-1/piwi* mutants, which have a defective piRNA pathway, most of the 23H-RNA loci had higher levels of 22G-RNAs in our sRNA-seq libraries, possibly an artifact due to loss of piRNAs and other 22G-RNAs (Figure 5C; Supplemental Table S3) (Batista et al. 2008; Das et al. 2008; Wang and Reinke 2008). Only *jmjc-1* had substantially reduced levels of 22G-RNAs in *prg-1* mutants, indicating that piRNAs do not have a major role in triggering 22G-RNA production from 23H-RNA loci (Figure 5C; Supplemental Table S3). In *ergo-1* mutants, which lose oogenic/embryonic 26G-RNAs, ∼50% of 23H-RNA loci were depleted of 22G-RNAs (Figure 5D; Supplemental Table S3) (Han et al. 2009; Gent et al. 2010; Vasale et al. 2010; Fischer et al. 2011). This was not unexpected as several of the 23H-RNA loci were previously classified as ERGO-1 class 26G-RNA targets (Han et al. 2009; Gent et al. 2010; Vasale et al. 2010).

23H-RNAs were not depleted in *mut-14 smut-1* mutants, indicating that the *Mutator* pathway involved in WAGO class 22G-RNA formation is not required for 23H-RNA formation (Supplemental Figure S5A; Supplemental Table S3). Consistent with our earlier conclusion that RDE-1 is not generally required for 23H-RNA formation or stability, only 5 individual 23H-RNAs were depleted in *rde-1* mutants (Supplemental Figure S5B; Supplemental Table S3). None of the 23H-RNAs were significantly depleted in *prg-1* and only 2 were weakly depleted in *ergo-1* mutants, indicating that the loss of 22G-RNA we observed in these mutants, particularly *ergo-1*, is not due to loss of 23H-RNAs (Supplemental Figures S5C-S5D; Supplemental Table S3). Our results demonstrate that multiple small RNA pathways, most notably the 23H-RNA, ERGO-1 class 26G-RNA, and WAGO class 22G-RNA pathways converge on a shared set of targets (Figure 5E).

### Fertility and fitness of *rde-1* and 23H-RNA mutants

We next assessed whether 23H-RNAs might have a role in health or fitness by examining the morphology, developmental timing, behavior, and fertility of *rde-1* mutants. We also generated a full deletion allele (*ram37*) of the locus producing the most abundant 23H-RNAs, F43E2.6, to assess whether individual 23H-RNA loci are important for development. We did not observe any obvious developmental defects or a difference in developmental timing in *rde-1* or F43E2.6 mutants (Figures 6A-6B). Consistent with a previous study, we observed a small reduction in fertility in *rde-1(ne219)* mutants at 25°C, which was reproducible in separate experiments (Figure 6C; Supplemental Figure S6A) (Seroussi et al. 2023). However, we did not observe a significant difference in a distinct loss-of-function allele of *rde-1*, *ne300*, although there was a strong tendency for both *rde-1* alleles to produce fewer progeny (Figure 6C; Supplemental Figure S6A). Nor did we observe a consistent reduction in fertility in *rde-1* mutants grown at the more favorable temperature of 20°C (Supplemental Figure S6B). Fertility was also not affected in F43E2.6 mutants (Figure 6C). The *rde-1(ne219)* mutant, with an amino acid substitution, has a weaker RNAi-defective phenotype than the *rde-1(ne300)* mutant, which has a premature stop codon (Tabara et al. 1999b; Watts et al. 2020). Perhaps the stronger phenotype in *rde-1(ne219)* mutants is due to background mutations, or it may produce a mildly toxic RDE-1 protein, which we did not explore.

**Figure 6.**
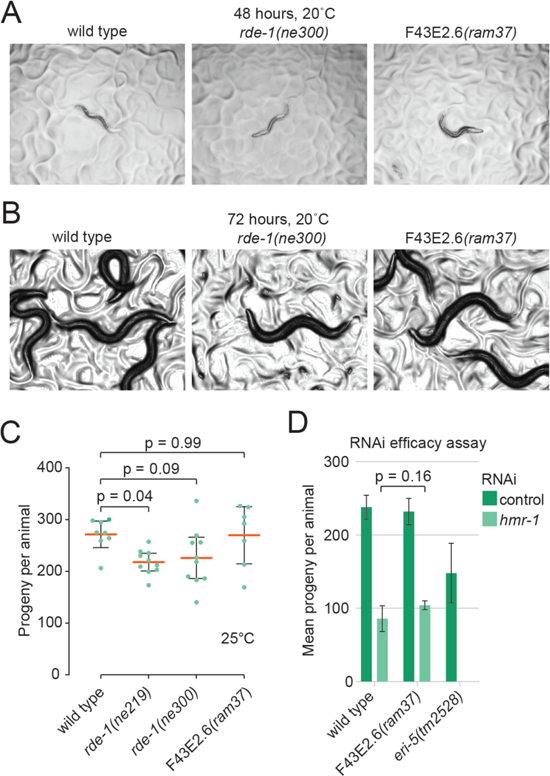
*rde-1* loss-of-function mutants are healthy. (A) Representative images of L4 stage larvae for each of the genotypes indicated. Animals were grown for 48 hours at 2O°C. (B) Representative images of adult stage animals for each of the genotypes indicated. Animals were grown for 72 hours at 2O°C. (C) Each data point is the number of progeny produced by a single animal for each of the genotypes indicated. The orange bars show the means and the error bars show the 95% confidence intervals. n=7-10 animals per genotype. One-way ANOVA followed by a Dunnett test was used for statistical analysis. (D) RNAi-effi-ciency assay. Mean progeny per animal for each of the genotypes indicated is shown following control or *hmr-1* RNAi. Error bars are SD between 3 biological replicates. The p value was calculated using a two-sample t-test.

We then did mobility assays to identify any behavioral abnormalities in *rde-1* and F43E2.6 mutants. The overall speed, numbers of reversals, and number of turns were not significantly different between wild type and each of the mutants (Supplemental Figures S6C-S6E). Collectively, these results suggest that neither F43E2.6 or RDE-1, and thus presumably the 23H-RNA pathway, is essential for general health under ideal laboratory conditions.

Perhaps loss of abundant 23H-RNAs could free up bandwidth in the RNAi pathway and thereby enhance RNAi. However, loss of F43E2.6 did not enhance RNAi against *hmr-1*, which in wild type animals has a modest response to dsRNA feeding that leads to reduced fertility, but which can be increased in mutants with enhanced RNAi efficiency, such as *eri-5* (Figure 6D) (Fischer et al. 2008; Thivierge et al. 2012). It is possible that additional loss of abundant 23H-RNAs could trigger an enhanced RNAi phenotype, although it is not known at which steps the RNAi machinery is limiting but it may be downstream of RDE-1 (Billi et al. 2014).

## DISCUSSION

In this study we characterized the endogenous canonical siRNAs of *C. elegans*. We predicted that such siRNAs were largely missed in earlier studies because of their relatively low abundance compared to other classes of small RNAs. Indeed, our analysis uncovered several hundred new small RNAs with the hallmarks of canonical siRNAs from RDE-1 co-IPs, most of which were quite rare and often undetectable in cell lysates. Our results add to the already expansive repertoire of *C. elegans* small RNAs and demonstrate that endogenous siRNAs like those found in mice and flies are prevalent, albeit typically at low levels, in worms. We compiled these siRNAs, including their sequence variants, along with miRNAs, 21U-RNAs, 22G-RNAs, and 26G-RNAs into a GFF3-formatted annotation file compatible with the *C. elegans* genome releases WS235-WS290+ and WS279 annotations, which can be utilized in most high-throughput sequencing data analysis software. For greatest sensitivity, we recommend RNA from embryos or RDE-1 co-IPs is used when analyzing 23H-RNAs.

RNA editing is antagonistic to RNAi and related pathways, likely because mispairs introduced by ADARs inhibit dsRNA processing by Dicer (Scadden and Smith 2001; Knight and Bass 2002; Tonkin and Bass 2003; Yang et al. 2005; Ohta et al. 2008; Heale et al. 2009; Sebastiani et al. 2009; Wu et al. 2011; Warf et al. 2012; Reich et al. 2018; Fischer and Ruvkun 2020). It therefore stands to reason that siRNAs would be upregulated in animals deficient in RNA editing activity. Our discovery pipeline was designed to identify siRNAs produced in effectively wild type animals and that associate with the RNAi Argonaute RDE-1.

Presumably a similar analysis in ADAR mutants would uncover many additional endogenous siRNAs, however, the biological relevance of siRNAs produced only in ADAR mutants is questionable outside of an RNA editing context. In contrast, the siRNAs characterized in this study may have biological roles in wild type animals, for example, in response to viral infection or stress, and will likely serve as useful readout for defects in RNAi, negating the need for exogenous dsRNA treatment when characterizing mutant alleles implicated in small RNA pathways.

The mild phenotypes we observed in *rde-1* mutants suggests that under optimal growth conditions, 23H-RNAs are not particularly important for development. Additionally, any reduction in fitness in *rde-1* mutants could relate to altered miRNA activity rather than to a role in the 23H-RNA pathway. The lack of a clear phenotype in *rde-1* mutants is reminiscent of the mild phenotypes originally observed in Drosophila AGO2 mutants, which appear superficially wild type but have non-essential roles in germ cell development and male fertility, presumably due to loss of siRNA activity (Okamura et al. 2004; Xu et al. 2004; Deshpande et al. 2005; Wen et al. 2015). Similar subtle phenotypes could emerge for *rde-1* mutants through more detailed analyses.

The hyper-enrichment of 23H-RNAs and exogenous siRNAs in RDE-1 co-IPs over most miRNAs may relate to the double stranded nature of the duplex intermediates. In support of this, and consistent with earlier findings, nearly all of the most highly enriched miRNAs in our GFP::RDE-1 co-IPs derive from near-perfect duplexes (<3 mismatches) (Seroussi et al. 2023). Whether this relates to the duplex itself or to biogenesis or sorting factors, such as RDE-4, a dsRNA-binding protein specific to 26G-RNA and siRNA production, is unclear (Billi et al. 2014). It is also possible that RDE-1 binds small RNA duplexes produced by Dicer indiscriminately, but only retains a subset of siRNAs, for example, those for which it can cleave and discard the passenger strand, such as might occur in duplexes with near perfect complementarity (Steiner et al. 2009).

Future studies exploring 23H-RNAs will likely uncover biological roles for these small RNAs. For now, several knowledge gaps exist. For example, do 23H-RNAs act in cis or trans? Do they have roles in preventing transposon mobilization or silencing endogenous genes? Are their roles masked because they act upstream of a transgenerational silencing pathway that persists in the absence of the 23H-RNA triggers? Additionally, it will be important to explore their biogenesis and the mechanism of target gene silencing. Our study providing a framework for addressing these questions and for further exploration of the 23H-RNA pathway.

## METHODS

### Strains

USC1080[*rde-1(rde-1(cmp133[(gfp + loxP + 3xFLAG)::rde-1])* V], WM27[*rde-1(ne219)* V], and WM45[*rde-1(ne300)* V] were previously described (Tabara et al. 1999b; Svendsen et al. 2019). The TM134[F43E2.6*(ram37)* II] strain was generated using CRISRP-Cas9-mediated genome editing with purified Cas9 protein and synthesized guide RNAs (CCUCCAACGAUCCACCACAG and AAUUCUUGAUAAAGUCCAGG) in wild type (N2) *C. elegans*, resulting in a replacement of a 2,483 bp sequence from positions −819 to 1664 in relation to the start of the coding sequence with the sequence TTA (Integrated DNA Technologies) (Gasiunas et al. 2012; Jinek et al. 2012; Cong et al. 2013; Mali et al. 2013). Analysis of other strains in this study relied on published datasets.

### RNAi

Synchronized L1 larvae were placed on *E. coli* HT115 expressing dsRNA matching *nrfl-1* and *oma-1*, *hmr-1*, *dcr-1*, or empty vector (L4440), and were grown at 20°C (Kamath et al. 2003). Gravid adults were collected for RNA isolation or GFP::RDE-1 co-IP at 72 hours. The *pos-1* RNAi data was from a previous experiment (Svendsen et al. 2019).

### Protein-small RNA co-IPs

USC1080[*rde-1(cmp133[(gfp + loxP + 3xFLAG)::rde-1])* V] animals were grown on RNAi treatment for 72 hours after L1 synchronization. GFP::FLAG::RDE-1 was co-IP’d from two biological replicate samples with ∼21,000 animals each. Animals were flash-frozen in liquid nitrogen and ground in in 50 mM Tris-Cl, pH 7.4, 100 mM KCl, 2.5 mM MgCl2, 0.1% Igepal CA-630, 0.5 mM PMSF, and 1X Pierce Protease Inhibitor Tablets (Pierce Biotechnology, cat# 88266). Cell lysates were cleared by centrifugation for 10 min at 12,000 RCF. Cleared lysates were split into input and co-IP fractions. The co-IP fractions were incubated with GFP-Trap Magnetic Agarose Beads (Proteintech, cat# gtma-100) for 1 hour. Beads were washed in lysis buffer and split into RNA and protein fractions. Western blot analysis was used to confirm enrichment of GFP::RDE-1 in co-IPs relative and wild type samples were used to confirm specificity of GFP antibodies for GFP::RDE-1.

### RNA isolation

RNA was isolated from whole wild type and mutant animals and from cleared cell lysates or GFP::RDE-1 co-IPs using Trizol followed by two rounds of chloroform extraction according to the manufacturer’s recommendations (Life Technologies, cat# 15596018). RNA was precipitated in isopropanol overnight at −80°C in the presence of 20 ug glycogen.

### Small RNA sequencing

For wild type, *rde-1(ne219)*, GFP::RDE-1 co-IP, and cell lysate libraries, ∼16-30-nt RNA was size selected on 17% polyacrylamide/urea gels from RNA pre-treated with RppH (Almeida et al. 2019). Small RNA sequencing libraries were prepared using the NEBNext Multiplex Small RNA Library Prep Set for Illumina following the manufacturer’s recommendations with the 3’ ligation step changed to 16°C for 18 hours to improve capture of methylated small RNAs (New England Biolabs, cat# E7300S). Small RNA PCR amplicons were size selected on 10% polyacrylamide non-denaturing gels. All other RNA sequencing was previously described (Welker et al. 2010; Rybak-Wolf et al. 2014; Svendsen et al. 2019; Reed et al. 2020; Montgomery et al. 2021; Seroussi et al. 2023).

### Data analysis

sRNA-seq data was analyzed and plotted using the default configuration in tinyRNA (Tate et al. 2023). Reference genome sequences and annotations were from the *C. elegans* genome WS279 release (Davis et al. 2022). Custom Python scripts, freely available on request, were used to identify complementary sequences with 2-nt 3’-overhangs enriched in GFP::RDE-1 co-IPs from tinyRNA alignment tables. 23H-RNA and other small RNA annotations are available at MontgomeryLab.org. Hairpin structures were predicted and drawn with RNAfold (Gruber et al. 2008). Microsoft Excel, GraphPad Prism, R, IGV, and Illustrator were also used for plotting and statistical analysis of data (Robinson et al. 2011).

### Quantitative real-time PCR

Quantitative real-time-PCR of small RNAs was done with TaqMan reagents and a custom probe targeting an F43E2.6 siRNA (sequence: UUUGCCGAUGUUUCUGAGAUGUC), and miR-1 (Life Technologies, assay name: hsa-miR-1, cat# 4427975) following the manufacturer’s recommendations (Life Technologies, cat# 4331348). F43E2.6 siRNA and miR-1 Ct values were assayed using a CF96 Real-Time PCR Detection System (Bio-Rad). Ct values were averaged across three technical replicates for each of 3 biological replicates.

Mean F43E2.6 siRNAs levels relative to miR-1 were calculated using the 2-ddCt method (Livak and Schmittgen 2001). Two-sample t-tests were used to compare differences between conditions.

### Brood size assays

To assess general fertility, the parental generation of the animals in which fertility was assayed was grown at 20°C until the L4 stage and then transferred to the experimental temperature (20°C or 25°C). Individual animals were grown on OP50 in individual wells of 24 well plates starting at the L4 stage and transferred to new wells every 24 hours. The number of progeny in each well was counted the following day. The process was repeated until the cessation of egg laying. To assess fertility for the RNAi efficiency assay, 20 animals per replicate were grown on RNAi from L1 larval stage until day 2 of adulthood. The parental animals were then removed and the numbers of progeny was assessed by washing them off each plate ∼24 hours later into 2 ml of M9 and then averaging the number of larvae in 5 x 10 μl aliquots and extrapolating the total.

### Behavior assays

For each assay, five 1-day old hermaphrodites were transferred from 60mm NGM plates with OP50 to a 60mm chemotaxis plates (CTX) (5 mM KH2PO4/K2HPO4 [pH 6.0], 1 mM CaCl2, 1 mM MgSO4, 2% agar), without food and left to roam for 2 minutes to adjust to the new plate. The plate was then placed onto the Wormlab Imaging System (MBF bioscience) and animal movements were recorded for 5 minutes (Roussel et al. 2014). Tracks were manually controlled for head and tail orientation blinded to genotype. Worms that showed no movement for 5 minutes or that left the field of view within 1 minute were excluded from analysis. Absolute speed (µm/s), total number of turns, and total number of reversals were captured using the Wormlab imaging system and exported to GraphPad Prism for statistical analysis.

### Statistics

For sRNA-seq, the DESeq2 R package was used within the tinyRNA pipeline to compute normalized counts using the geometric mean method and perform statistical analysis using the Wald test (Love et al. 2014; Tate et al. 2023). R and GraphPad Prism were used for ANOVA and TukeyHSD or Dunnett tests, and Microsoft Excel was used to calculate t-tests where indicated.

## DATA ACCESS

All raw and processed sequencing data generated in this study have been submitted to the NCBI Gene Expression Omnibus (GEO; https://www.ncbi.nlm.nih.gov/geo/) under accession number GSE254442.

## ACKNOWLEDGEMENTS

Thanks to Maritza Soto-Ojeda and Reese Sprister for help with media and solutions, and Josh Svendsen and Kristen Brown for help with small RNA-seq libraries. Strains were provided by the CGC, which is funded by the National Institutes of Health (P40 OD010440). This work was supported by the National Institutes of Health [R35GM119775 to T.A.M. and R01NS115947 to F.J.H.].

---

**Supplement Figure S1.**
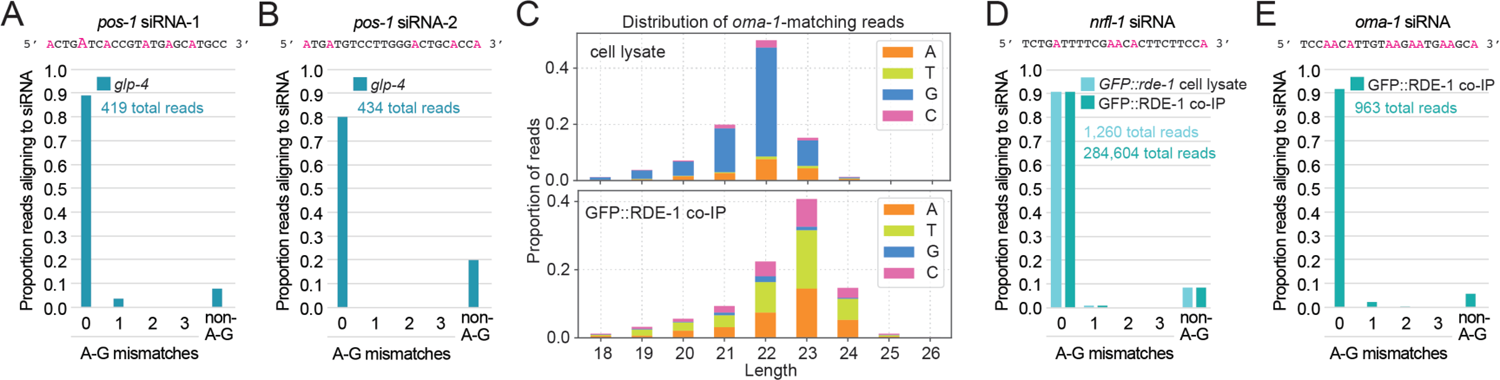
RNA editing and RDE-1 association with exogenous Γ siRNAs. (A-B) The proportion of small RNA-seq reads from two different *pos-1* siRNAs with 0-3 A to G and total (1-3) non A to G mismatches to the genome in *glp-4(bn2)* mutants undergoing *pos-1* RNAi. Adenosines (As) are colored magenta and their heights are roughly proportional the frequency of A to G editing. n=1 biological replicate. Gravid adult stage animals grown at 25°C. (C) Size and 5’-nt distribution of *oma-1* siRNAs from cell lysates and GFP::RDE-1 co-IP libraries following *nrfl-1* and *oma-1* RNAi. n=2 biological replicates. Data from one representative library shown. Gravid adult stage animals.(D-E) The proportions of reads from a *nrfl-1* siRNA (D) or an *oma-1* siRNA (E) with mismatches to the genome, as in A-B, in cell lysates and GFP::RDE-1 co-IPs from animals undergoing *nrfl-1* and *oma-1* RNAi. n=2 biological replicates. Data from one representative library shown.

**Supplemental Figure S2.**
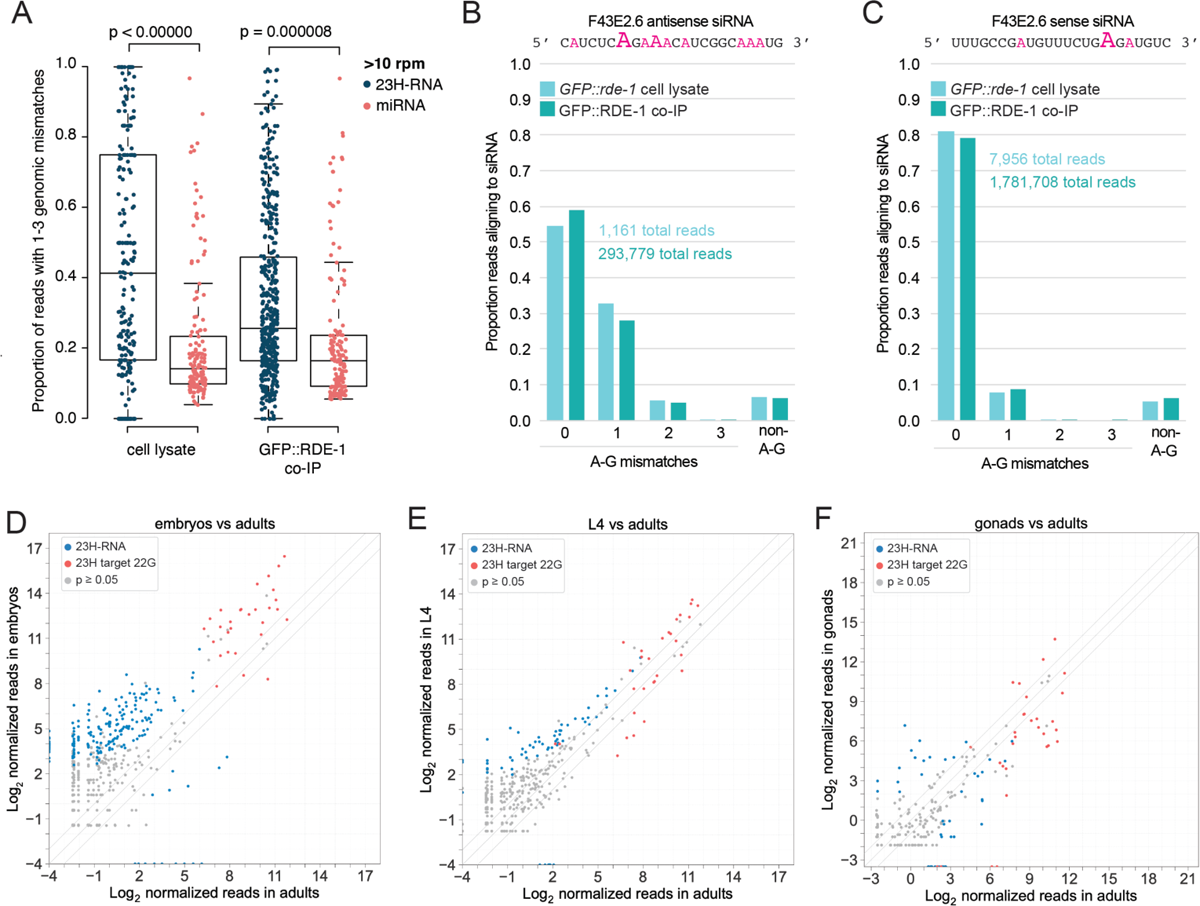
RNA editing and expression of endogenous siRNAs. (A) Each data point in the box plot shows the proportion of reads in a 23H-RNA or miRNA with 1-3 mismatches to the genome in small RNA-seq libraries from cell lysates and GFP::RDE-1 co-IPs. Only small RNAs with an average of >10 reads per million total in GFP::RDE-1 co-IPs are shown. n=2 biological replicates. P values were calculated using a TukeyHSD test. (B-C) The proportion of small RNA-seq reads from two different F43E2.6 siRNAs with 0-3 A to G and total (1-3) non A to G mismatches to the genome. Adenosines (As) are colored magenta and their heights are roughly proportional the frequency of A to I editing. 1 of 2 biological replicates is shown. (D-F) Each data point in the scatter plots shows the average log_2_ geometic mean-normalized small RNA-seq reads in gravid adult C. *elegans* on the x axis and embryos (D), L4 stage larvae (E), or dissected distal gonads (F) on the y axis. n=3 biological replicates for each developmental stage. Samples from different experiments.

**Supplemental Figure S3.**
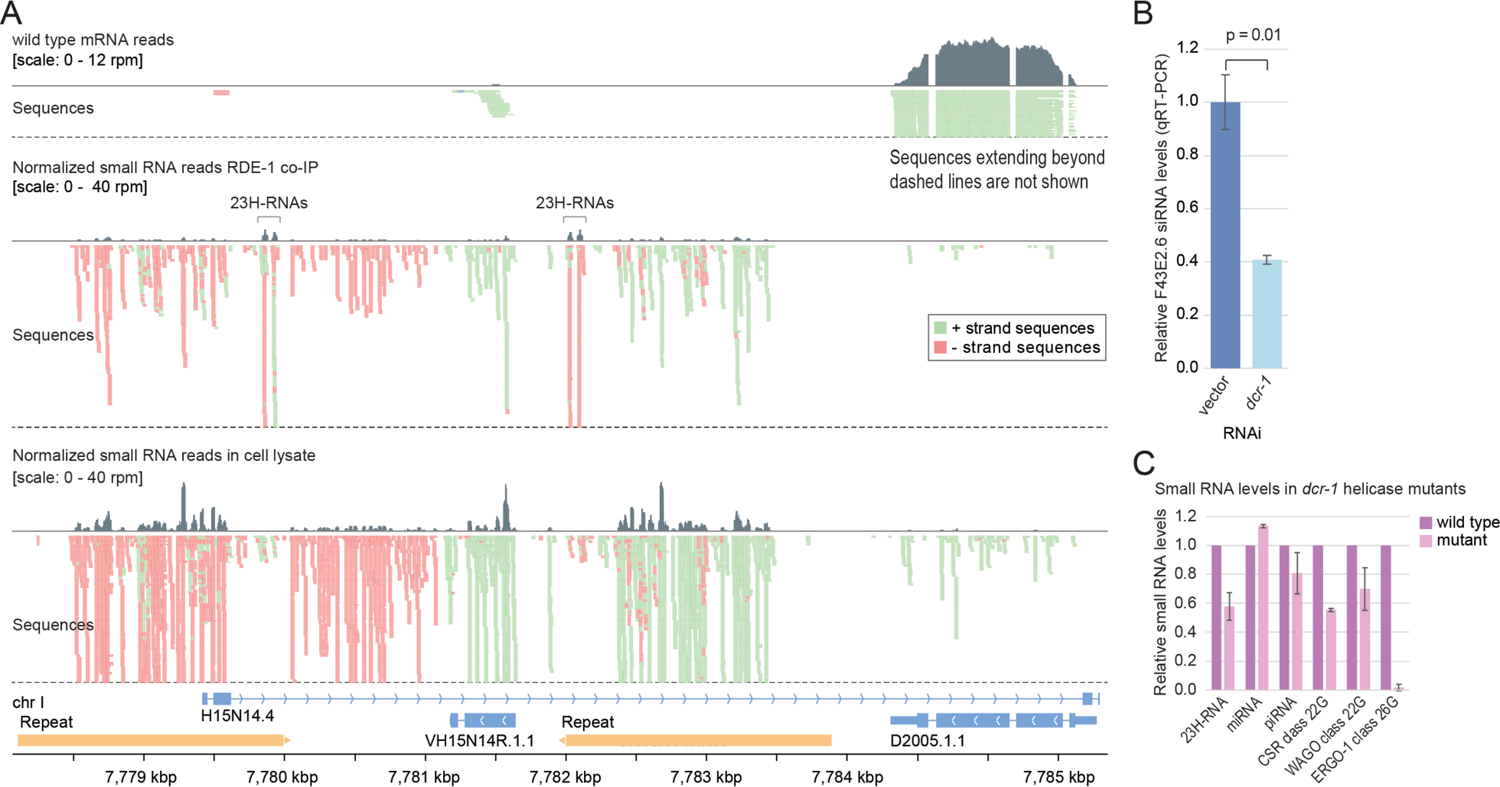
23H-RNA distribution across an inverted repeat and dependence on Dicer. (A) mRNA reads from wild type animals and small RNA-seq reads from cell lysates and GFP::RDE-1 co-IPs plotted across an inverted repeat 23H-RNA locus. Brackets indicate 23H-RNA clusters. 1 of 2 biological replicates is shown. (B) Relative F43E2.6 siRNA levels after control (L4440) or *dcr-1* RNAi as determined by qRT-PCR. miR-1 levels were used for normalization. Error bars are SD between 3 biological replicates. The p value was calculated using a two-sample t-test. (C) Relative levels of the various classes of small RNAs in *dcr-1* mutants rescued with wild type *dcr-1* ora *dcr-1* helicase mutant. n=1 for wild type *dcr-1* rescue and n=2 for *dcr-1* helicase mutant rescue. Error bars for rescue with the *dcr-1* helicase mutant are SD. Mixed stage animals.

**Supplemental Figure S4.**
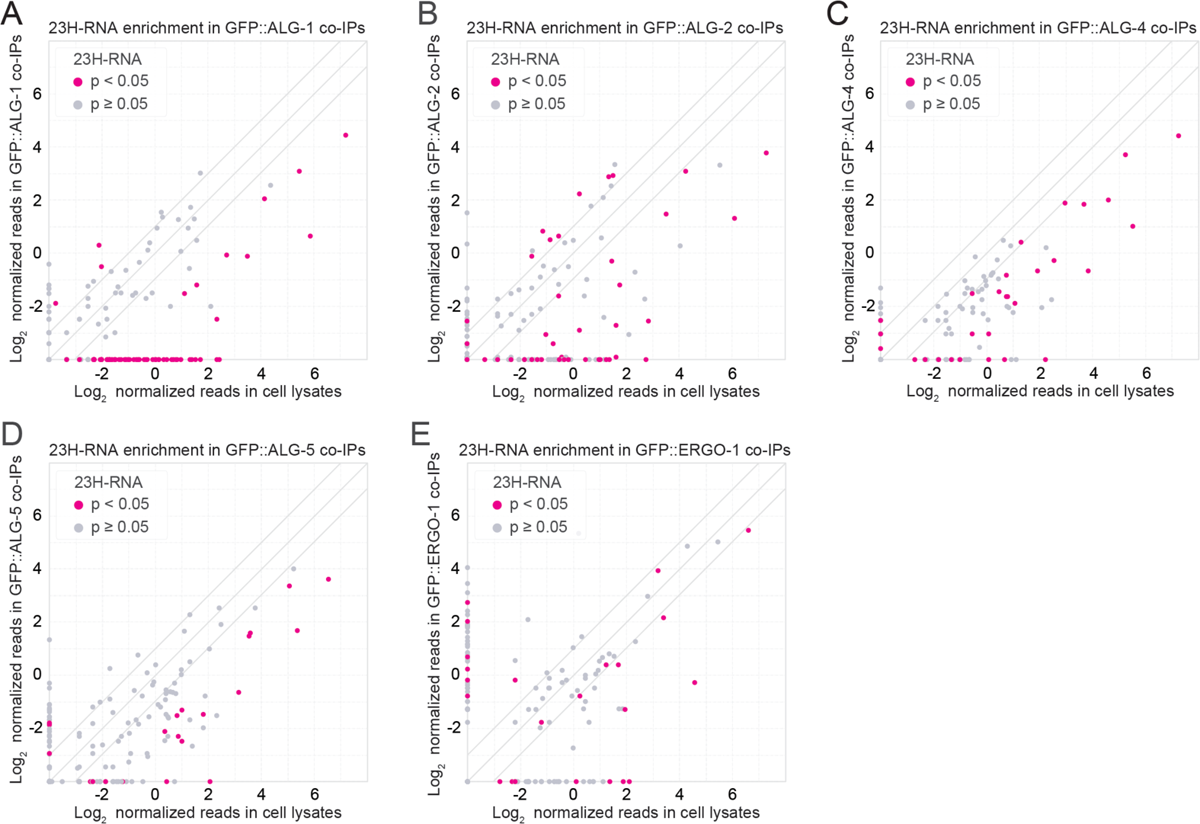
23H-RNA enrichment or depletion in Argonaute co-IPs. (A-E) Each data point in the scatter plots shows the average log_2_ rpm-normalized small RNA-seq 23H-RNA reads from cell lysates on the x axis and Argonaute co-IPs on the y axis. (A) GFP::ALG-1 co-IPs. (B) GFP::ALG-2 co-IPs. (C) GFP::ALG-4 co-IPs, L4 larvae. (D) GFP::ALG-5 co-IPs. (E) GFP::ERGO-1 co-IPs. n=2 biological replicates for each condition. Adult stage animals except for GFP::ALG-4 co-IPs and cell lysates which were L4 larval stage.

**Supplemental Figure S5.**
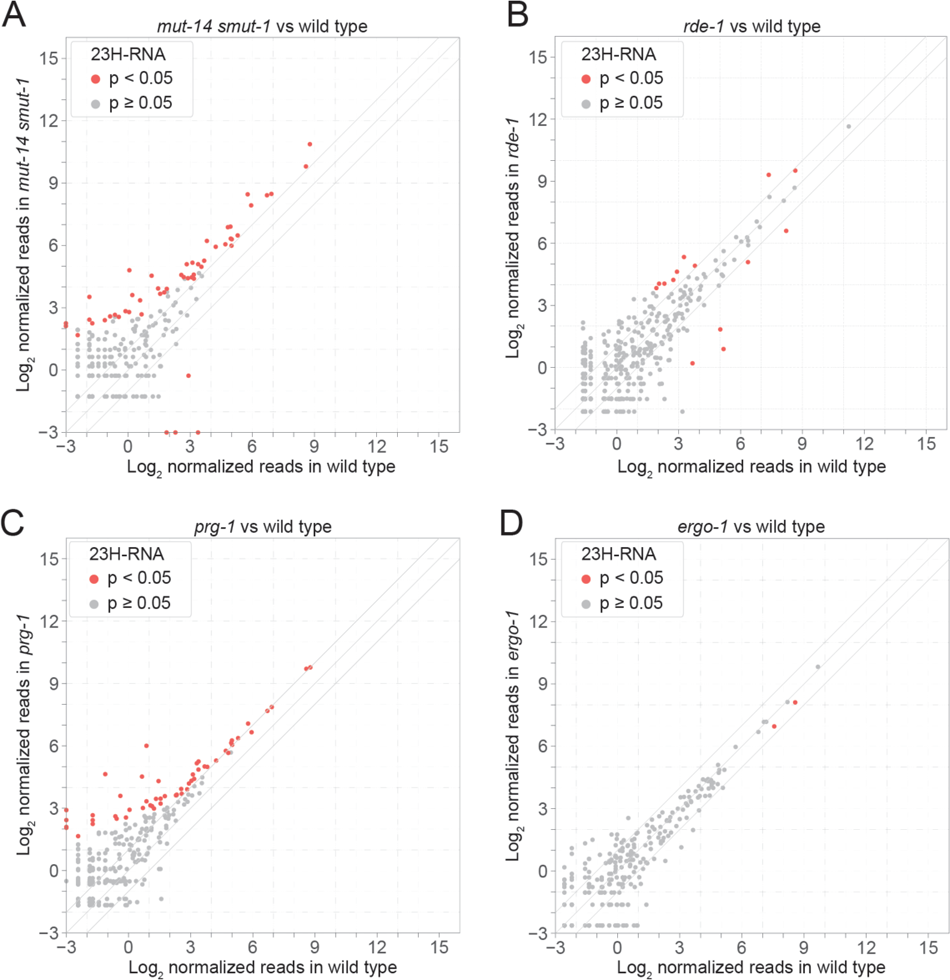
Genetic requirements for 23H-RNAs. **(A-D)** Each data point in the scatter plots shows the average log_2_ geometic mean-normalized small RNA-seq 23H-RNA reads in wild type *C. elegans* on the x axis and *mut-14(mg461) smut-1(tm1301) (A), rde-1(ne219) (B), prg-1(n4357)* (C), or *ergo-1(tm1860)* (D) on the y axis. Gravid adult stage animals. 3 biological replicates each, except *mut-14(mg461) smut-1(tm1301)* which had only 2.

**Supplemental Figure S6.**
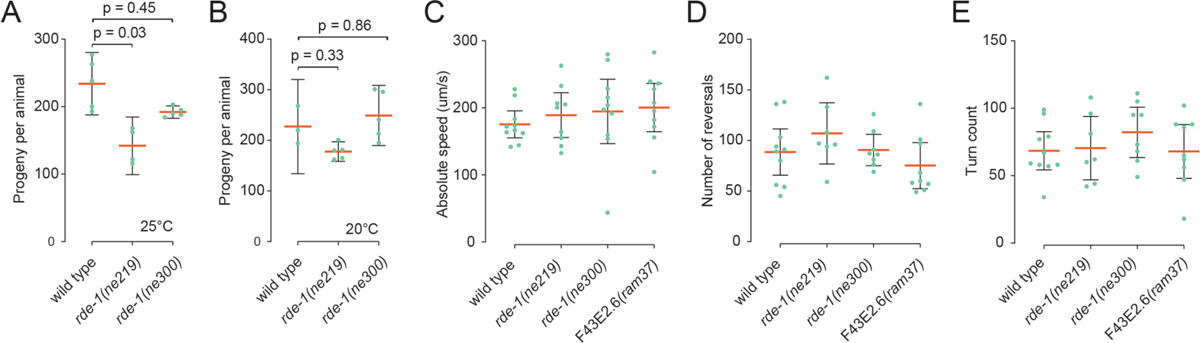
Fertility and behavior of *rde-1* and F43E2.6 mutants. (A) Each data point is the number of progeny produced by a single animal for each of the genotypes indicated grown at 25°. The orange bars show the means and the error bars show the 95% confidence intervals. n=4-5 animals per genotype. (B) As in A but for animals grown at 2O°C. n=3-4 animals per genotype. (C) Absolute speed (forward + reverse animal body movement). (D) Number of reversals. (E) Number of turns. For C-E, n=9-10 animals per measurement. One of three independent experiments, each with ∼10 animals per replicate, is shown. Measurement were taken over 5 minutes. All statistical analysis was done using one-way ANOVA followed by Dunnett tests.

